# Hemoglobin alpha regulates T-lymphocyte activation and mitochondrial function

**DOI:** 10.1101/2025.08.01.668160

**Authors:** Emily C. Reed, Tatlock H. Lauten, Tamara Natour, Lauren J. Pitts, Caroline N. Jojo, Brooke L. Griffin, Sreeram Pasupuleti, Adam J. Case

## Abstract

We have recently discovered hemoglobin alpha a1 (Hbα-a1 mRNA and Hbα protein) in T-lymphocytes and previously reported that its expression was sensitive to mitochondrial redox perturbations. However, outside of its occurrence and basic characterization, the functional role of Hbα in T-lymphocytes remained unknown. Herein, we identify Hbα in both CD4^+^ and CD8^+^ T-lymphocyte subsets, and found its expression is highly dynamic, differs between the two subtypes, and is dependent upon activation stage. Further, the loss of Hbα by use of a novel T-lymphocyte-specific Hbα knock-out mouse impairs mitochondrial function, dysregulates cytokine production, and lowers the activation threshold primarily in CD4^+^ T-lymphocytes, indicating a critical role for Hbα within this subset. While these data suggested the loss of Hbα in T-lymphocytes may promote aberrant activation of autoreactive T-lymphocytes, surprisingly, we discovered that mice lacking Hbα in T-lymphocytes exhibited reduced severity of experimental autoimmune encephalomyelitis (EAE) compared to wild-type control animals. Interestingly, T-lymphocytes lacking Hbα *in vivo* appeared to function identically to wild-type controls, which did not explain the protection against EAE. In contrast, T-lymphocyte Hbα knock-out mice displayed significantly reduced levels of circulating immunoglobulins and CD40L expression compared to their wild-type counterparts during EAE, suggesting possible impaired intercellular communication. These data elucidate a previously unrecognized role for Hbα in T-lymphocyte function, which may have implications for hemoglobin-related diseases (i.e., hemoglobinopathies).

## Introduction

Over the past 30 years, research has challenged the dogma that hemoglobin is strictly confined to erythrocytes by identifying hemoglobin subunits expressed in a variety of other cell types, such as neurons (1–3), macrophages (4), mesangial cells (5), and others (6, 7). Through these studies, hemoglobin has been revealed to possess diverse redox abilities, including facilitating O_2_ or nitric oxide (NO) exchange, modulating iron utilization, antioxidant capabilities, protection during hypoxia, and mediating mitochondrial bioenergetics. In a preliminary report, we discovered hemoglobin alpha-a1 (Hbα-a1 mRNA and Hbα protein) expression in both mouse and human T-lymphocytes, showed its differential levels in various T-lymphocyte polarized states, and elucidated its increased production in response to redox perturbations, indicating a potential antioxidant function in these cells (8). We further demonstrated that Hbα overexpression led to an increase in mitochondrial membrane potential, suggesting that Hbα in T-lymphocytes may also play a role in mitochondrial homeostasis. Last, initial studies using T-lymphocyte-specific Hbα-a1 knock out mice (HbKO) depicted a significant decrease in the percentage of naïve CD4^+^ T-lymphocytes after psychological trauma, proposing irregular cellular activation in the absence of Hbα-a1. Altogether, these data suggest that Hbα may be necessary to prevent aberrant activation of T-lymphocytes by modulating the mitochondrial redox environment within these cells.

Given these interesting preliminary results demonstrating a functional role for hemoglobin in T-lymphocytes, we set out to explore if this protein was important in the regulation of autoimmunity. Herein, we identify unique temporal expression patterns of Hbα-a1 mRNA and Hbα protein in both CD4^+^ and CD8^+^ T-lymphocytes, with the loss of Hbα in T-lymphocytes exhibiting a decrease in mitochondrial metabolism, increase in proinflammatory cytokine production after 24-hour activation, and a decreased threshold for activation compared to wild-type (WT) T-lymphocytes. Incongruously, HbKO animals showed better disease outcomes in a preclinical model of autoimmune multiple sclerosis, known as experimental autoimmune encephalomyelitis (EAE), despite virtually identical T-lymphocyte inflammatory phenotypes in HbKO and WT animals. However, we uncover that this unexpected result may be due to impaired T-lymphocyte interactions with other immune cells, such as B-lymphocytes, hindering EAE disease development. Together, this work uncovers a vital functional role for Hbα in T-lymphocytes, and may have implications for hemoglobinopathies.

## Materials and Methods

### Mice

Wild-type C57BL/6J (#000664; shorthand WT) and B6.Cg-Tg(Cd4-cre)1Cwi/BfluJ (#022071; shorthand CD4-cre) mice were obtained from Jackson Laboratories (Bar Harbor, ME, USA). Conditional Hbα-a1 knock-out mice were graciously provided by Dr. Brant Isakson as previously described (9). CD4-cre is activated during the double positive developmental stage in the thymus leading to cre recombination in both mature CD4^+^ and CD8^+^ T-lymphocytes (10), therefore, conditional Hbα-a1 knock-out mice were crossed with CD4-Cre mice to generate pan-T-lymphocyte-specific modified progeny (HbKO). All mice were bred in house to eliminate shipping stress and microbiome shifts, as well as co-housed with their littermates (≤5 mice per cage) prior to the start of experimentation to eliminate social isolation stress. Mice were housed with standard pine chip bedding, paper nesting material, and given access to standard chow (#8604 Teklad rodent diet, Inotiv, West Lafayette, IN, USA) and water ad libitum. Male and female experimental mice between the ages of 8-14 weeks were utilized in all experiments. If no significant sex differences were observed, data are presented as pooled independent of sex to increase N while reducing unnecessary animal use (11). Experimental mice were randomized, and when possible, experimenters were blinded to the respective cohorts until the completion of the study. Mice were sacrificed by pentobarbital overdose (150 mg/kg, Fatal Plus, Vortech Pharmaceuticals, Dearborn, MI, USA) administered intraperitoneally. All mice were sacrificed between 7:00 and 9:00 Central Time to eliminate circadian rhythm effects on T-lymphocyte function. All procedures were reviewed and approved by Texas A&M University Institutional Animal Care and Use Committees.

### Mouse T-lymphocyte isolation, culture, and activation

Primary immune cells were collected from mice as previously described (8). Briefly, spleens or inguinal lymph nodes were collected and disrupted into a single cell suspension then passed through a 70 μM nylon mesh filter (#22363548, ThermoFisher Scientific). Erythrocytes were removed using red blood cell lysis buffer (150 mM NH_4_Cl, 10 mM KHCO_3_, 0.1 mM EDTA). Splenic T-lymphocytes were negatively selected using EasySep Mouse total, CD4^+^, or CD8^+^ T-cell negative magnetic isolation kit (StemCell Technologies #19851, #19852, #19853), per manufacturer’s instructions. T-lymphocytes were counted, and viability was assessed using Trypan Blue exclusion on a Bio-Rad TC20 Automated Cell Counter. For activation, cells were plated at 800,000 cells/mL with anti-CD3/28 Dynabeads (Dynabeads, #11456D) in a 1:1 cell to bead ratio in T-lymphocyte media consisting of RPMI media supplemented with 10% Fetal Bovine Serum, 2 mM Glutamax, 10 mM HEPES, 100 U/mL penicillin/streptomycin, and 50 μM of 2-mercaptoethanol. Cells were cultured in 5% CO_2_, 37**°**C incubator (HERAcell Vios 160i CO_2_ incubator, ThermoFisher Scientific).

### Experimental autoimmune encephalomyelitis (EAE) and restimulation

#### Disease induction

EAE was induced using the MOG_35-55_/CFA Emulsion kit (#EK-2110, Hooke Laboratories) as described in (12). Briefly, 9-14 week old mice were anesthetized and injected with 100 µl of myelin oligodendrocyte glycoprotein (MOG) emulsion in two separate subcutaneous locations. Two hours later, mice were anesthetized again and injected intraperitoneally with 100 µL of 100 ng/µL pertussis toxin in PBS, followed by a second dose after 24 hours. Beginning seven days later, mice were weighed and scored every day to assess disease progression. Disease scores (0–5) were based off a standardized clinical signs and symptom rubric, described in detail in (12). Separate cohorts of mice were sacrificed at either peak disease (day 14) or after disease relapse (day 28).

#### Restimulation

Total immune cells were isolated from spleens or inguinal lymph nodes as described above and cultured for 72 hours in the presence of 10 µg/mL of MOG_35-55_ (#NC0754019, Hooke Laboratories). Cells counts were assessed, and media was analyzed for extracellular proinflammatory cytokine production.

### Flow cytometry immunophenotyping and redox assessment

T-lymphocytes were immunophenotyped via 4-laser Attune NxT flow cytometer (ThermoFisher Scientific) as previously described (13). Cells were stained with 1:1000 dilutions of CD3ε PE-Cy7 (#25-0031-82, ThermoFisher Scientific), CD4 eFluor 506 (#69004182, ThermoFisher Scientific), CD8 Super Bright 702 (#67008182, ThermoFisher Scientific), CD19 APC-Cy7 (#BDB557655, Thermo Fisher Scientific), CD11b BV421 (#BDB562605, Thermo Fisher Scientific), CD11c APC, (#501129650, Thermo Fisher Scientific) antibodies along with 1 μM MitoSOX Red (#M36008, ThermoFisher Scientific) in RPMI media to assess mitochondrial reactive oxygen species (ROS) in various immune cell subpopulations. Mean fluorescence intensity (MFI) of MitoSOX Red was reported as a readout of mitochondrial ROS levels. Cells were stained with 100 nM tetramethylrhodamine ethyl ester (TMRE) (#T669, ThermoFisher Scientific) to measure mitochondrial membrane potential. Antibodies for staining other extracellular T-lymphocyte markers include CD28 APC (5014933, Thermo Fisher Scientific), CD27 Super Bright™ 600 (63027182, Thermo Fisher Scientific), and CD40L Alexa Fluor™ 488 (53154182, Thermo Fisher Scientific). Additionally, T-lymphocytes were also assessed for various polarization states, as previously described in (14, 15). Briefly, splenic or inguinal lymph node immune cells were treated with phorbol 12-myristate-13-acetate (10 ng/mL) (#5005820001, Thermo Fisher Scientific), BD GolgiPlug (#BDB555029, Thermo Fisher Scientific) and ionomycin (0.5 mg/mL) (#AAJ60628LB0, Thermo Fisher Scientific) for 4 hours at 37**°**C. Cells were then stained with Live-Dead Fixable Far Red viability dye (#501121530, Thermo Fisher Scientific) for 30 minutes, 4**°**C, washed, then followed by a 1:1000 dilution of extracellular antibodies for CD3ε PE-Cy7 (#25-0031-82, Thermo Fisher Scientific), CD4 eFluor 506 (#69004182, Thermo Fisher Scientific), and CD8 Alexa Fluor™ 488 (#69-0041-82, Thermo Fisher Scientific) for 30 minutes at 4**°**C. Afterwards, cells were fixed and permeabilized with FOXP3/Transcription Factor Staining Buffer Set according to manufacturer’s instructions (#501128857, Thermo Fisher Scientific). Cells were stained with 1:1000 dilutions of intracellular antibodies against IFNγ APC-eFluor780 (#47-7311-82, Thermo Fisher Scientific), IL-4 BV421 (#562915, BD Biosciences), FoxP3 PE-Cyanine5.5 (#35-5773-82, Thermo Fisher Scientific), and IL-17 BV711 (#407-7177-82, Thermo Fisher Scientific). All flow cytometry data was analyzed using FlowJo v10 software (BD Biosciences).

### RNA extraction, cDNA production, and quantitative real-time RT-PCR

T-lymphocyte RNA isolation and mRNA levels were assessed as previously described (8, 16). Briefly, mRNA was extracted using the RNAeasy plus mini kit (#74136, Qiagen) and quantified using NanoDrop One Spectrophotometer (#13400518, Thermo Scientific). RNA was then transformed into cDNA using ThermoFisher High-Capacity cDNA Reverse Transcriptase Kit (#4374967, Applied BioSystem). Generated cDNA was used for real time quantitative PCR. Primers for genes of interest were designed using NIH primer-BLAST spanning exon-exon junctions. Cq values were determined, and relative gene expression was calculated by comparing housekeeping 18s ribosomal gene expression to gene of interest (2^-ddCq^).

### Protein analysis

Protein was isolated from T-lymphocytes using RIPA lysis buffer (#PI89900, Thermo Scientific) and 1% Halt protease inhibitor cocktail (#PI87785, Thermo Scientific). Samples were subsequently subjected to sonication and centrifugation to obtain soluble protein, which was quantified using Pierce BCA protein assay kit (#PI23227, Thermo Scientific). Hbα protein was assessed via Total Protein Jess Automated Western Blot (Bio-techne) as described in (8, 16) using anti-Hbα primary antibody - Rabbit (#PIPA579347, Fisher) 1:20. Analysis was performed using Compass Software for Simple Western v6.1.0.

### Seahorse Mitochondrial Stress Test

T-lymphocyte mitochondrial metabolism assessment was performed as previously described (8, 16). Briefly, activated T-lymphocytes were plated in Seahorse XF RPMI media (#103576-100, Agilent) supplemented with 10 mM Seahorse XF Glucose (#103577-100, Agilent), 1 mM Seahorse XF Pyruvate (#103578-100, Agilent), 2 mM Seahorse XF L-Glutamine (#103579-100, Agilent). Cells were adhered to seahorse cell microplates using 1 µg/cm^2^ Cell-Tak (#354240, Corning) and seeded at a density of 200,000 cells per well. Mitochondrial inhibitors (1 µM Oligomycin, 1 µM 4-trifluoromethoxy-phenylhydrazone (FCCP), 0.5 µM Rotenone and Antimycin, #103015-100, Agilent), were injected into each well and oxygen consumption rate (OCR) was measured via Seahorse XFe96 Analyzer.

### Single-cell energetic metabolism by profiling translation inhibition (SCENITH)

Protein translation inhibition was measured as a proxy for ATP production, as described in detail in (17). Naïve or 72-hour activated splenic T-lymphocytes were plated (in duplicate per treatment group) at 500,000 cells per well in a 96 well plate. Cells were treated with 100 mM of 2-deoxy-D-glucose (2-DG), 1µM oligomycin (#103020-100, Agilent), or both for 45 minutes at 37**°**C. During the last 15 minutes of incubation, 10 µg/mL of puromycin (#A1113803, Thermo Fisher Scientific) was added to each well. Cells were washed, stained with Fixable Viability dye eFluor (#501128817, Thermo Fisher Scientific) for 30 minutes at 4**°**C, then subsequently stained with 1:1000 CD4 eFluor 506 (#69004182, ThermoFisher Scientific) and CD8 Alexa Fluor™ 488 (#69-0041-82, Thermo Fisher Scientific) extracellular antibodies. Cells were fixed and permeabilized (#501128857, Thermo Fisher Scientific), then intracellularly stained with 1:1000 anti-puromycin Alexa Fluor™ 647 antibody (#ab322729, Abcam) for 30 minutes at 4**°**C. Cells were then resuspended in PBS and fluorescence was detected using the 4-laser Attune NxT flow cytometer (ThermoFisher Scientific). MFI of puromycin in live CD4^+^ and CD8^+^ T-lymphocytes was calculated using FlowJo v10 software (BD Biosciences). Average MFI of each experimental group for each sample was calculated, and mitochondrial dependence, glycolytic capacity, glucose dependence, and fatty acid/amino acid oxidation capacity were determined as described in (17).

### Cytokine protein analysis

Cytokine protein levels from *in-vivo* (i.e., plasma) and *ex-vivo* (i.e., media normalized to cell counts) experiments were analyzed using Mesoscale multi-plex technology (Mesoscale Discovery), as described in (14). Briefly, protein concentrations were assessed using custom Mesoscale Discovery U-Plex kits (Mesoscale Discovery, Rockville, MD, USA) simultaneously targeting IL-1β, IL-2, IL-4, IL-6, IL-10, IL-17A, IFNγ, and TNFα per manufacturer’s instructions. Analysis was performed using a Mesoscale QuickPlex SQ 120 and analyzed using Mesoscale Discovery software.

### Immunoglobulin ELISA

Plasma immunoglobin concentrations for IgG1, IgG2a, IgG2b, IgG2c, IgA, IgM, lambda light chain, and kappa light chain immunoglobulins were determined using Invitrogen™ Mouse Ig Isotyping Instant ELISA™ Kit (#501125163, Thermo Fisher). Plasma was collected from EAE animals as described above and diluted 1:10 prior to analysis. Assay was performed according to manufacturer’s instructions with analysis performed at 450 nm using SpectraMax i3x multimode plate reader (Molecular Devices).

### Complete blood count (CBC)

Blood parameters were assessed by complete blood count. Mice were sacrificed and blood was obtained via cardiac puncture. Undiluted blood (25 µL) was then transferred into a microvette (#NC9141704, Fisher Scientific) and analyzed by Abaxis VetScan HM5 Color Hematology System.

### Activation threshold

T-lymphocyte activation threshold was assessed by isolating CD4^+^ and CD8^+^ T-lymphocytes as described above, and plating with constant concentration of anti-CD28 (1 µg/mL) (#76556-222, VWR) and variable concentration of anti-CD3 (0.01-10 µg/mL) (#76556-606, VWR) soluble antibodies. Media was harvested 24 hours later, and cytokine protein expression was assessed via Mesoscale.

### Migration assay

T-lymphocyte migration was assessed using a transwell migration assay. Splenic CD4^+^ and CD8^+^ T-lymphocytes were isolated and activated with Dynabeads for 72 hours. Cells were then counted and 1 x 10^6^ cells were plated into the apical side of a 5 μM Millicell^®^ 24 well hanging cell culture insert (#PTMP24H48, Millipore Sigma) in FBS-free media. FBS-free media only (control) or media supplemented with 100 ng/mL CXCL12 (#478-MR-025/CF, Biotechne) or 100 ng/mL RANTES (#460-SD-050/CF, Biotechne) was added on the basolateral side of the insert. After 4 hours, basolateral side media was collected and migrated T-lymphocyte counts were assessed by flow cytometry. Ratio of migrated cells was calculated by dividing chemokine migrated cell count by control migrated cell count.

### Mitochondrial DNA and mass assessment

DNA from naïve and 72-hour activated T-lymphocytes was isolated using GeneJet Genomic DNA Purification Kit (#K0722, Thermo Fisher Scientific). Mitochondrial (*Nd1* and *Cytb*) and nuclear (*Gusb* and *B2m*) DNA content was assessed via qPCR. Mitochondrial mass was quantified in naïve and 72-hour activated T-lymphocytes using MFI of MitoTracker Green FM (#M46750, Thermo Fisher Scientific) by flow cytometry.

### Statistical analysis

All data presented as mean ± standard error of the mean (SEM), with N values representing individual mice listed in figure legends. Normality was assessed using Shapiro-Wilk normality test before statistical analysis. For two group comparisons, Student’s t-test was utilized. For experiments with 3 or more groups, an ordinary one-way ANOVA was performed. Experiments containing two categorical groups were assessed using two-way ANOVA or mixed-effects analysis. All statistics were completed using GraphPad Prism version 10.5.0.

## Results

### Activation state of T-lymphocytes determines intracellular Hbα levels and impact on mitochondria

While we reported the expression of Hbα-a1 in T-lymphocytes (8), the dynamics of its expression during cellular activation remained unknown. To address this knowledge gap, splenic CD4^+^ and CD8^+^ T-lymphocytes were isolated from WT mice, Hbα-a1 mRNA and Hbα protein levels were assessed 24-, 48- and 72-hours post-activation, and values normalized to the respective naïve cell-type levels. In contrast to significant reductions in mRNA levels at 24 hours post-activation, Hbα protein content increased in both CD4^+^ and CD8^+^ T-lymphocytes compared to naïve T-lymphocytes (**Figure 1A-D**). Congruent with 48-hour mRNA levels, Hbα protein decreased in both subtypes, but diverged between CD4^+^ and CD8^+^ at 72 hours (**Figure 1A-D**). The highly variable expression of Hbα throughout the first 72 hours of T-lymphocyte activation highlights the temporal and diverse regulation of Hbα within activated T-lymphocytes, and suggests differential importance within T-lymphocyte subtypes.

**Figure 1:**
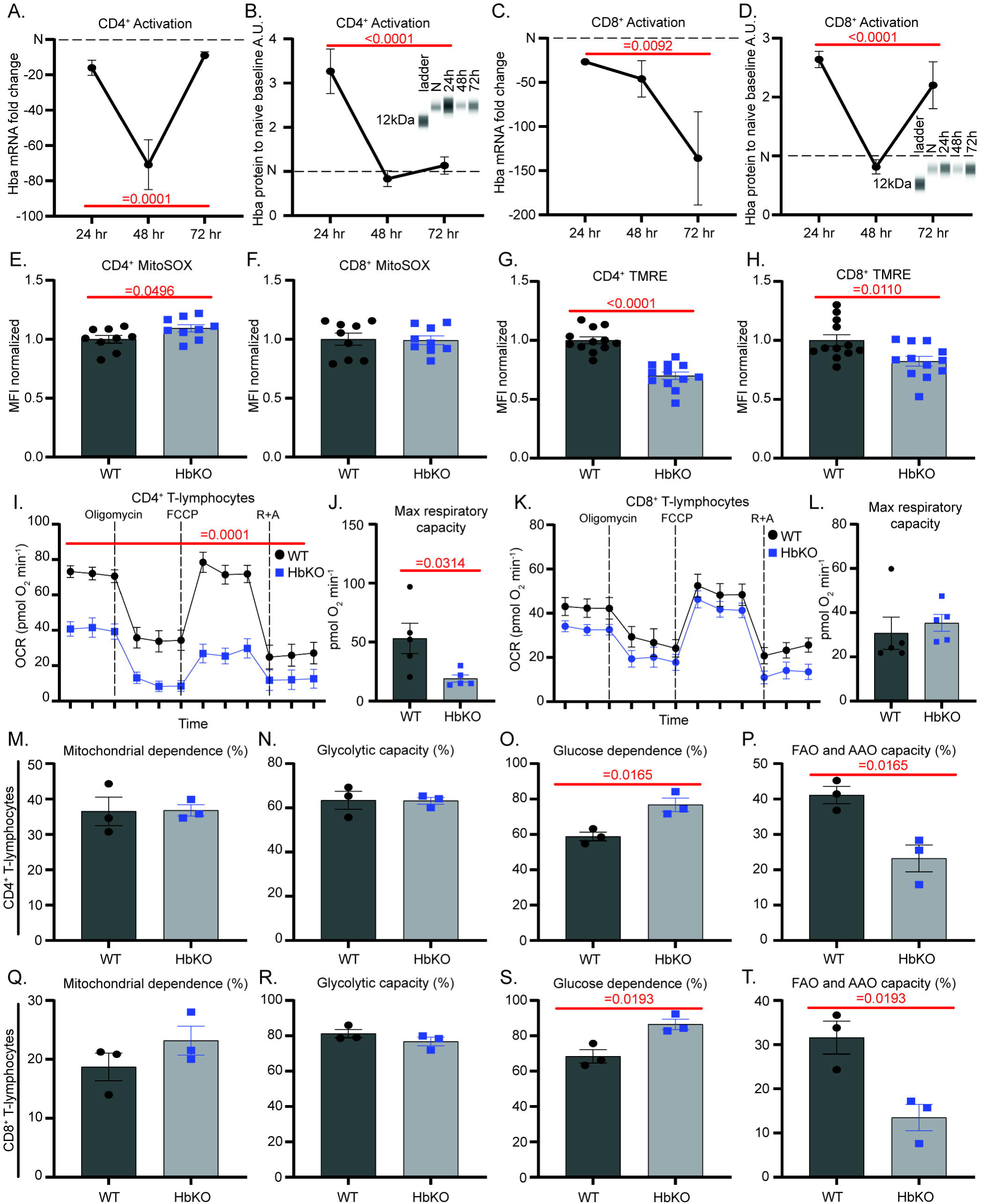
Activation state of T-lymphocytes determines intracellular Hbα levels and impact on mitochondria. **A-D**: Hbα-a1 mRNA (N=7) and Hbα protein expression (N=7) over 24, 48, 72-hour time points after activation of CD4^+^ (**A-B**) and CD8^+^ (**C-D**) T-lymphocytes compared to naive T-lymphocytes (N=6). **E-H**: Splenic CD4^+^ and CD8^+^ T-lymphocytes were isolated and activated for 72 hours, then assessed by flow cytometric analysis of MitoSOX (**E-F**) and TMRE MFI (**G-H**). **I-L**: Splenic CD4^+^ and CD8^+^ T-lymphocytes were isolated and activated for 72 hours, then assessed using Seahorse mitochondrial stress test (N=5). R+A = rotenone + antimycin A. **M-T**: Splenic T-lymphocytes were activated for 72 hours, then puromycin MFI was measured via SCENITH protocol in CD4^+^ and CD8^+^ T-lymphocytes to obtain calculations graphed (reported in percentage). Statistics measured by a one-way ANOVA with Dunnett’s multiple comparisons test, unpaired Student’s t-test, or two-way ANOVA with Šídák’s multiple comparisons test where appropriate.

To investigate the mechanistic function of Hbα-a1 in T-lymphocytes, we next generated a T-lymphocyte specific Hbα-a1 knock out (HbKO) mouse model (**Supplementary Figure 1A**). Cre-mediated DNA recombination was confirmed to be confined solely to T-lymphocytes (**Supplementary Figure 1B**), and T-lymphocyte-specific loss of Hbα-a1 did not impact any erythrocyte or leukocyte parameter on complete blood count analysis (**Supplementary Figure 1C-F**). Further, Hbα-a1 knock-out did not affect the percentage or cell counts of either CD4^+^ or CD8^+^ splenic T-lymphocytes, suggesting loss of Hbα-a1 does not significantly impact T-lymphocyte viability during development (**Supplementary Figure 1G-J**). *Ex-vivo* proliferative capacity was also not hindered by the loss of Hbα-a1 in either T-lymphocyte subtype (**Supplementary Figure 1K-L**).

Given that we previously demonstrated that T-lymphocyte Hbα-a1 expression was highly sensitive to redox perturbations primarily stemming from the mitochondria (8), we next sought to explore T-lymphocytes mitochondrial bioenergetics in the absence of Hbα-a1. We observed a modest, but significant, increase in mitochondrial reactive oxygen species (ROS) levels only in activated CD4^+^ HbKO T-lymphocytes compared to WT CD4^+^ cells (**Figure 1E**), with no differences observed in activated CD8^+^ or naïve cells of either subtype (**Figure 1F, Supplemental Figure 1M-N**). Additionally, no significant differences were noted in either total cellular ROS or NO levels in either CD4^+^ and CD8^+^ HbKO T-lymphocytes (**Supplemental Figure 1O-R**), which suggested a primary impact of Hbα on the mitochondria. Indeed, mitochondrial membrane potential of both CD4^+^ and CD8^+^ HbKO activated T-lymphocytes was significantly decreased compared to their WT counterparts **(Figure 1G-H),** indicating the potential of mitochondrial metabolic perturbations. Furthermore, CD4^+^ HbKO T-lymphocytes displayed significantly lower mitochondrial O_2_ consumption, with the cells showing a significantly lower maximum respiratory capacity compared to WT CD4^+^ activated T-lymphocytes (**Figure 1I-J**). However, while CD8^+^ HbKO T-lymphocytes showed a slight decrease in O_2_ consumption, these effects did not reach statistical significance **(Figure 1K-L)**. Additionally, protein translation is known to be tightly coupled to ATP production, and thus inhibition of ATP synthesis pathways (i.e., glycolysis and or mitochondrial oxidative phosphorylation) can provide an indirect snapshot of metabolic dependencies (17). Interestingly, neither CD4^+^ or CD8^+^ activated HbKO T-lymphocytes showed any differences in mitochondrial dependence or glycolytic capacity compared to WT T-lymphocytes **(Figures 1M-N, 1Q-R)**. However, there was a significant increase in glucose dependence, coupled with a significant decrease in fatty acid/amino acid oxidation in HbKO T-lymphocytes **(Figures 1O-P, 1S-T)**. Interestingly, these mitochondrial changes with the loss of Hbα-a1 are likely not due differences in mitochondrial number or mass, due to no significant differences in mitochondrial DNA content or MitoTracker green MFI in either subtype or activation state (Supplementary Figure 2A-J).

Altogether, these data suggest Hbα-a1 plays a significant role in maintaining mitochondrial function and redox balance in activated T-lymphocytes, particularly in CD4^+^ subtypes.

### Activated HbKO T-lymphocytes exhibit an accelerated and enhanced proinflammatory phenotype

We and others have previously demonstrated an important role for mitochondrial metabolism and redox in shaping T-lymphocyte activation and effector functions (13, 18–21). Given the impact of Hbα on mitochondrial redox and metabolic parameters, we hypothesized Hbα loss would significantly alter activation in T-lymphocytes. Strikingly, CD4^+^ HbKO T-lymphocytes produced higher levels of all cytokines post-activation compared to WT T-lymphocytes, suggesting the Hbα knock-out leads to a loss of cytokine regulation post-activation **(Figure 2A-D, Supplementary Figure 3A-D)**. In contrast, only IL-6 and IL-17A were impacted in CD8^+^ T-lymphocytes (**Figure 2E-H, Supplementary Figure 3E-H**), once again supporting a greater impact of Hbα loss in CD4^+^ T-lymphocytes. Furthermore, HbKO CD4^+^ T-lymphocytes displayed higher levels of cytokines throughout an increasing amount of CD3 stimulation, with HbKO CD8^+^ T-lymphocytes showing minimal impact (**Figure 2 I-P**), suggesting a lower threshold of activation specifically in CD4^+^ cells. Overall, these data indicate that the loss of HbKO in T-lymphocytes potentiates the magnitude and timing of activation-associated cytokines, which primarily impacts CD4^+^ subtypes.

**Figure 2:**
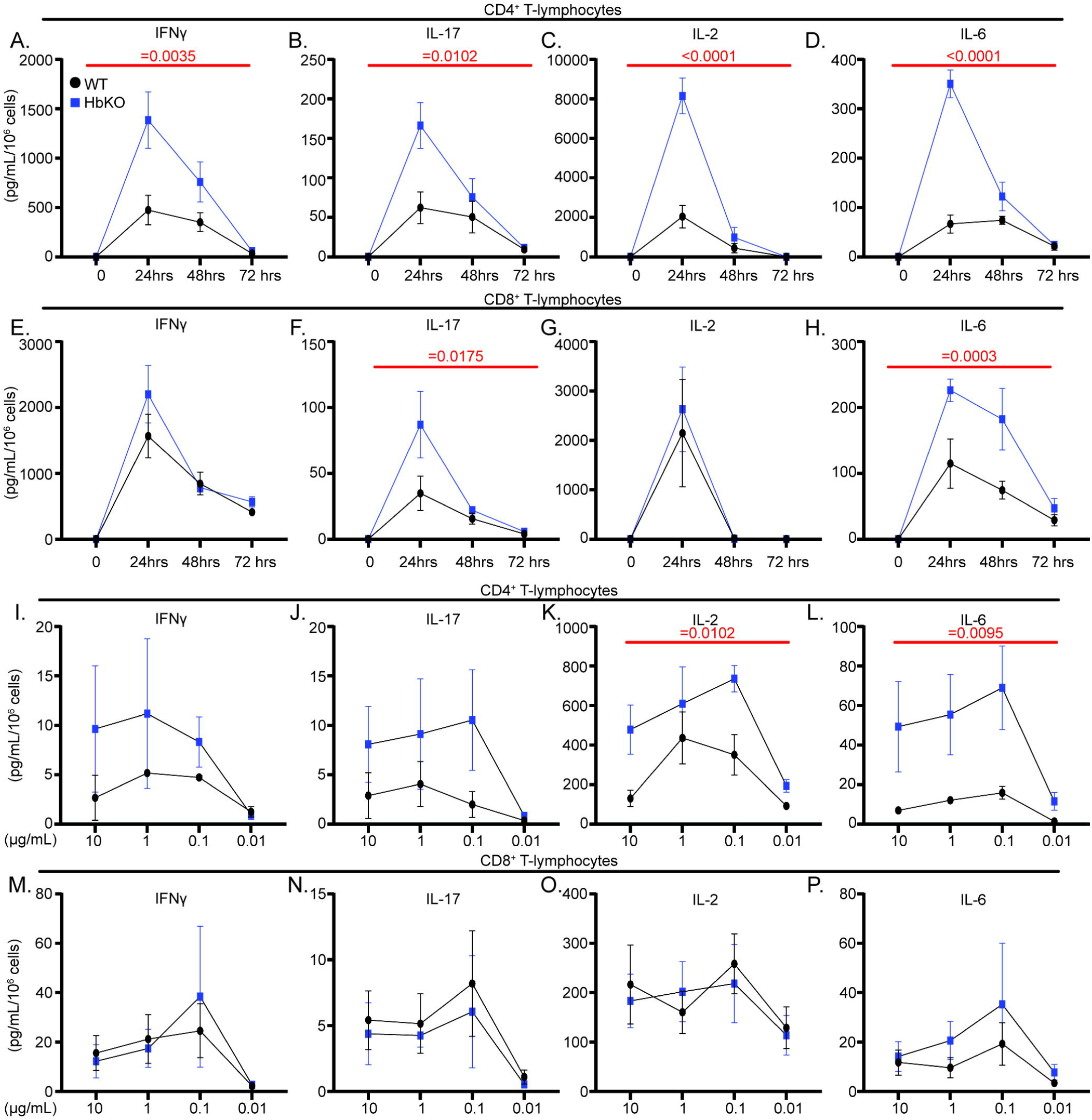
Activated HbKO T-lymphocytes exhibit an accelerated and enhanced proinflammatory phenotype. **A-H**: Extracellular cytokine protein measured from CD4^+^ (**A-D**) and CD8^+^ (**E-H**) T-lymphocytes at 24, 48, 72 hours post-activation (pg/mL per 10^6^ cells) (N=5). **I-P**: Extracellular cytokine protein measured from CD4^+^ (**I-L**) and CD8^+^ (**M-P**) T-lymphocytes treated for 24 hours with anti-CD28 and varying concentrations of anti-CD3 (N=3). Statistics measured using unpaired Student’s t-test, mixed-effects analysis, or two-way ANOVA with Šídák’s multiple comparisons test where appropriate.

### HbKO EAE animals exhibit better disease phenotypes but similar inflammatory profiles as WT EAE animals

To assess the impact of Hbα loss on T-lymphocyte function *in vivo*, we induced a T-lymphocyte-dependent model of experimental autoimmune encephalomyelitis (EAE) in WT and HbKO animals. Given the observation of potentiated cytokine production in the absence of Hbα *ex vivo*, we hypothesized that HbKO animals would have significantly worsened EAE disease progression. Contrary to our hypothesis, HbKO animals had delayed disease progression, significantly lower EAE disease severity scores, and diminished weight loss over the experimental period **(Figure 3B-C, Supplementary Figure 4B-C)**. Surprisingly, circulating plasma cytokines levels were identical between WT and HbKO animals at both 14 and 28 days post immunization (**Figure 3D-G, Supplementary Figure 4D-G).** Additionally, the only change in T-lymphocyte population noted was a significantly decreased proportion of CD4^+^ T-lymphocytes in draining lymph nodes at 28 days in HbKO animals (**Figure 3I**). In contrast, all other populations (including T_H_1 and T_H_17 cells, which are highly causal in EAE disease progression (22)) at 14 and 28 days in both the spleen and lymph nodes showed no differences compared to WT animals (**Figure 3H-O, Supplementary Figure 4H-O, 5A-D),** emphasizing that loss of T-lymphocyte Hbα does not inhibit *in vivo* CD4^+^ polarization and likely does not account for the difference in EAE scores. Moreover, there was no significant difference in percentage of other splenic immune cell populations, total immune cell infiltration into the spinal cord, or splenic T-lymphocyte mitochondrial ROS between the two genotypes **(Supplementary Figure 5E-K)**. Last, T-lymphocyte memory also appeared unimpacted by the loss of Hbα given that MOG_35-55_ antigen recall showed no differences in cellular proliferation or cytokine production per cell at either 14 or 28 days post-EAE immunization in the spleen (**Figure 3P-S, Supplementary Figure 3P-S, Supplementary Figure 5L**) or lymph nodes (data not shown).Together, these puzzling findings show that HbKO mice possess a significantly decreased susceptibility to EAE compared to WT mice, but the mechanism underlying this difference does not appear to be directly related to primary T-lymphocyte effector functions.

**Figure 3:**
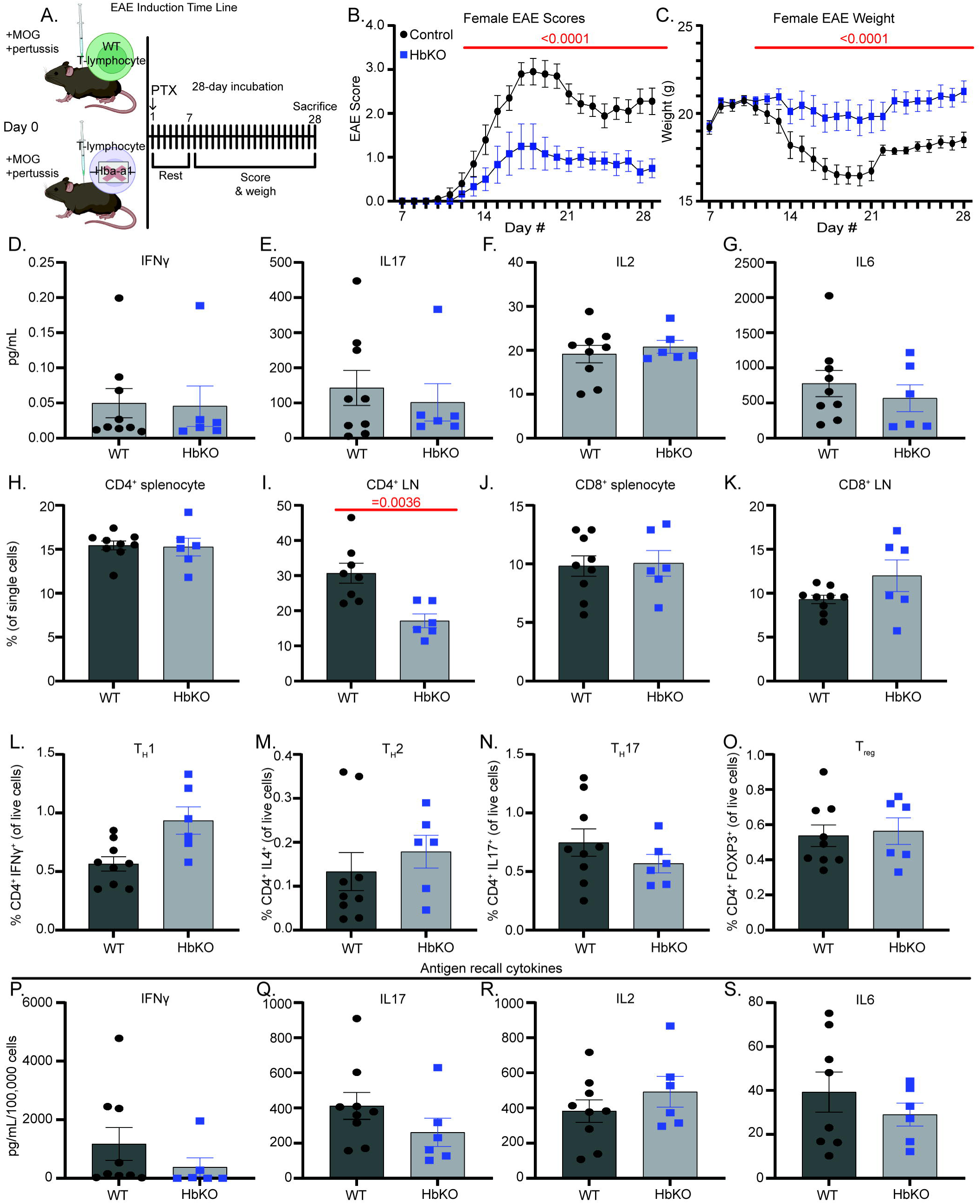
HbKO EAE animals exhibit better disease phenotypes but similar inflammatory profiles as WT EAE animals. **A**: Schematic of 28-day EAE experimental design. **B-C**: EAE severity scores (0–5) and weights (g) over 28-day incubation period (WT N=14, HbKO N=10). **D-G**: Protein concentration (pg/mL) of cytokines in plasma. **H-K:** Percentage of CD4^+^ and CD8^+^ present in the spleen and inguinal lymph nodes at day 28. **L-O**: Percentage of splenic CD4^+^ polarized T-lymphocytes. **P-S**: Splenocytes restimulated with 10 μg/mL MOG_35-55_ for 72 hours, then assessed for extracellular cytokine protein concentration (pg/mL per 10^6^ cells). Statistics measured by two-way ANOVA with Šídák’s multiple comparisons test or Student’s t-test where appropriate.

### Loss of Hbα impairs T-lymphocyte intercellular communication

Given the decreased percentage of CD4^+^ T-lymphocytes in the draining lymph nodes of HbKO EAE mice compared to WT mice (**Figure 3I**), we postulated that the loss of Hbα may alter the ability of T-lymphocytes to appropriately migrate to chemotactic stimuli. However, neither CD4^+^ or CD8^+^ HbKO T-lymphocytes demonstrated any deficiency in chemotactic migration *ex-vivo* (**Figure 4A-H**), suggesting this is likely not the mechanism of decreased EAE susceptibility with Hbα-a1 knock-out in T-lymphocytes.

**Figure 4:**
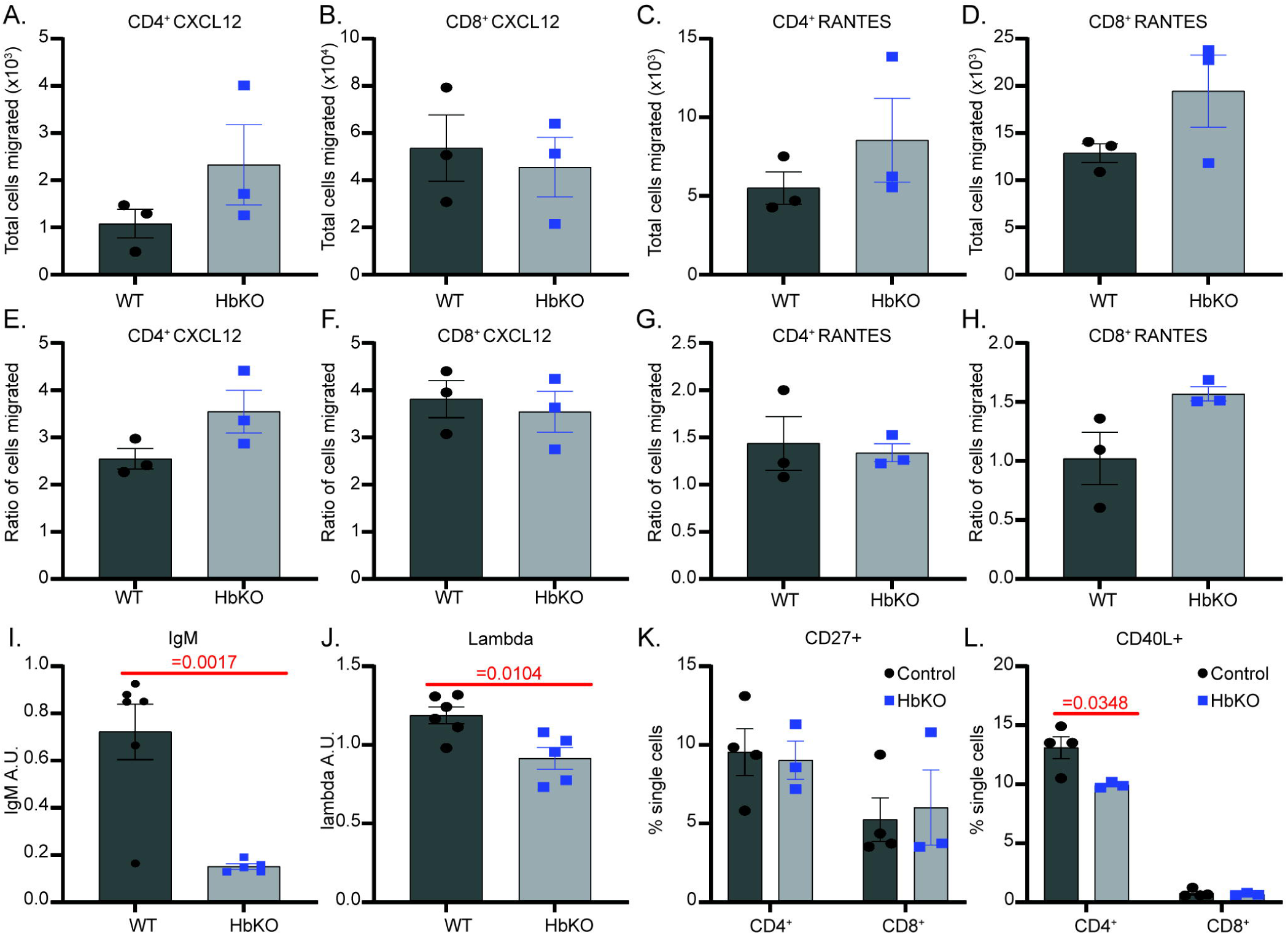
Loss of Hbα impairs T-lymphocyte intercellular communication. **A-H**: Splenic T-lymphocytes were isolated and activated with a 1:1 ratio of Dynabeads for 72 hours, replated into transwell inserts over control, RANTES, or CXCL12 media for 4 hours, then total cell counts (**A-D**) and ratio of cells migrated over control well migration (**E-H**) were assessed by flow cytometry. **I-J**: Relative IgM and lambda immunoglobulin concentrations in plasma from 14-day EAE animals (AU: arbitrary units). **K-L**: Splenic T-lymphocytes were isolated and assessed at 4-hour activated (**L**) and 24-hour activated (**K**) for receptor expression by flow cytometry. Statistics measured using Student’s t-test. Statistics (only significant shown) were measured by two-way ANOVA with Šídák’s multiple comparisons test or student’s t-test where appropriate.

Understanding that CD4^+^ T-lymphocytes interact with and potentiate inflammation from other immune cells (i.e., B-lymphocytes) to promote EAE disease progression, we next queried levels of circulating antibodies in the respective mouse models. Interestingly, while IgG and IgA antibodies remain unchanged, HbKO EAE animals exhibited significantly lower levels of plasma IgM and lambda immunoglobins compared to WT EAE animals, suggesting the loss of T-lymphocyte Hbα caused a significant impact on specific B-lymphocyte antibody output **(Figure 4I-J)**. To explore this further, we assessed receptors and ligands that are essential for the interaction between T-lymphocytes and B-lymphocytes. While we did not find any differences in the number of cells expressing the costimulatory receptor CD27 between HbKO and WT T-lymphocytes (**Supplementary Figure 5L**), we did observe significantly fewer CD4^+^ HbKO T-lymphocytes expressing CD40L (**Figure 4K**).

Congruent with previous findings, no changes were observed in CD8^+^ T-lymphocytes (**Figure 4K**). Together, these data promote the hypothesis that the loss of Hbα in T-lymphocytes leads to insufficient crosstalk with other immune cells, but additional work is warranted to explore the full depth and repertoire of these deficient interactions.

## Discussion

Hemoglobin is becoming increasingly recognized as a protein expressed in a variety of cell types beyond erythrocytes, and has been shown to possess varying redox functions and regulatory mechanisms throughout its diverse cellular potency (6, 7). We recently reported that Hbα is expressed in T-lymphocytes and its expression is malleable to redox perturbations, suggesting a redox regulatory role (8). Herein, we expand and refine our knowledge on Hbα expression at both the RNA and protein levels, explore its role in the context of T-lymphocyte metabolism and activation, and define differences between CD4^+^ and CD8^+^ T-lymphocyte subtypes. Altogether, our data suggest Hbα is an important T-lymphocyte redox-regulatory protein that is crucial for maintaining mitochondrial bioenergetics and proper T-lymphocyte activation, particularly in CD4^+^ T-lymphocytes.

Previously, we demonstrated that Hbα-a1 mRNA expression was rapidly down regulated within 24 hours in total T-lymphocytes after activation, likely due to activation-promoted epigenetic remodeling (8). Herein, we discovered that Hbα-a1 mRNA expression continues to decrease compared to naïve levels after 48 hours post activation in both CD4^+^ and CD8^+^ cells, but diverges depending on subtype at 72 hours.

Interestingly, Hbα protein expression revealed a differential pattern than the mRNA, with protein conversely increasing after 24 hours post activation compared to naïve T-lymphocytes. These disparate levels suggest the potential for a post-transcriptional regulatory mechanism between Hbα-a1 mRNA and Hbα protein. Moreover, this robust increase in Hbα protein at 24 hours may be in response to the massive burst of ROS produced during T-lymphocyte activation (18, 20, 23). While this burst of ROS is imperative for activation, uncontrolled ROS can lead to cell death, and is therefore tightly controlled by a subsequent increase of antioxidants, such as glutathione, Nrf2 activation, and as potentially described herein, Hbα (23, 24). In accordance with the mRNA expression, protein expression decreased in both subtypes after 48 hours, but once again increased after 72 hours activation. This increase in Hbα protein at the later time point may represent the early stages of polarization occurring in which certain subtypes of T-lymphocytes (i.e., memory T-lymphocytes, regulatory T-lymphocytes, etc.) display significantly elevated levels of Hbα as we have previously shown (8). Overall, Hbα seems to be intricately regulated at both the RNA and protein level in the major T-lymphocyte subtypes and activation states. This dynamic regulatory pattern suggests Hbα is not a passive bystander in T-lymphocytes, but instead plays an important and vital role that requires intricate and tight control.

While the functional role of non-canonical hemoglobin likely varies depending on cell type, it has been hypothesized that its function may be linked to mitochondrial energetics (25–30). We too recently reported that the loss of Hbα in T-lymphocytes significantly increased MitoSOX oxidation in a mouse model of psychological trauma (8). To further explore this phenomenon here, we examined the mitochondrial phenotype of naïve and activated T-lymphocyte subtypes. While there were no discernable differences between naïve WT and HbKO T-lymphocytes, HbKO T-lymphocytes activated for 72 hours possessed greater MitoSOX oxidation and significantly lower TMRE fluorescence, suggesting mitochondrial dysfunction. This dysfunction was more prominent in CD4^+^ T-lymphocytes, which concurrently showed a greater difference in mitochondrial bioenergetics compared to their WT counterparts. Additionally, both CD4^+^ and CD8^+^ activated HbKO T-lymphocytes showed a greater reliance on glucose as opposed to fatty acids/amino acids, further suggesting the loss of Hbα may greatly impact mitochondrial metabolism. Interestingly, the observed increase in MitoSOX oxidation, lower TMRE fluorescence, and decrease in overall OCR in HbKO CD4^+^ T-lymphocytes is very similar to T-lymphocytes that lack an important mitochondrial antioxidant, manganese superoxide dismutase (MnSOD), further suggesting Hbα may play a similar role in modulating the mitochondrial redox environment of activated T-lymphocytes (19).

In addition to disrupted mitochondrial bioenergetics, the loss of Hbα significantly altered the timing and magnitude of cytokine production after activation. Both CD4^+^ and CD8^+^ HbKO T-lymphocytes produced higher concentrations of specific cytokines such as IL-6 and IL-17A, but CD4^+^ T-lymphocytes showed a more pronounced phenotype with an apparent global loss of cytokine regulation, particularly 24 hours post-activation. Additionally, CD4^+^ HbKO T-lymphocytes produced higher concentrations of cytokines at 24 hours in response to lower concentrations of CD3 stimuli, suggesting that the loss of Hbα lowers the threshold of activation. The combination of greater mitochondrial dysfunction and increased production of proinflammatory cytokines in HbKO CD4^+^ T-lymphocytes may suggest that Hbα plays an important role in CD4^+^ T-lymphocyte activation and cytokine regulation.

Given both the highly proinflammatory phenotype and the decreased threshold of activation in HbKO T-lymphocytes *ex vivo,* we hypothesized that loss of Hbα would accelerate autoimmune risk or progression.

Autoimmune disorders are characterized by aberrant autoreactive immune cells, primarily of the adaptive immune system. Interestingly, the loss of hemoglobin has been hypothesized to correlate with an increased likelihood of developing autoimmune disorders, in a disease group broadly called hemoglobinopathies (31–35). Hemoglobinopathies are a class of genetic diseases in which one or more subunits of hemoglobin are impacted, and are one of the world’s most prominent inherited genetic disorders affecting around 7% of the human population (36). The consequences of hemoglobinopathies are primarily attributed to decreased oxygen (O_2_) delivery by erythrocytes, but it is unclear how mutations or alterations in hemoglobin would impact the development of autoimmune disorders. In fact, low O_2_ has been shown to decrease T-lymphocyte responses (37), which would suggest hemoglobinopathies would be more likely to suppress the prevalence of comorbid autoimmune diseases. Therefore, we posited that the loss or impact of hemoglobin mutations in T-lymphocytes may be partially explain the elevated risk of autoimmunity in hemoglobinopathies

EAE induction with MOG_35-55_ was chosen due to the important role of T-lymphocytes in developing the disease, as compared to MOG_1-125_ which is a primary B-lymphocyte-mediated response (38, 39). To our surprise and contrary to our hypothesis, HbKO animals fared better, showing delayed disease progression and less severe phenotypes. Intriguingly, both genotypes exhibited similar levels of circulating cytokines, splenic T-lymphocyte percentages, CD4^+^ T-lymphocyte polarization subtypes, and antigen recall responses, indicating that decreased disease severity was likely not due to decreased T-lymphocyte viability, differentiation, or cytokine production. HbKO animals did have less CD4^+^ T-lymphocytes present in the inguinal lymph nodes at 28 days compared to WT EAE lymph nodes, but this difference was not apparent at the height of disease severity (i.e., 14 days). Moreover, HbKO T-lymphocytes showed no differences in transwell migration towards either CXCL12 or RANTES (a cytokine known to recruit T-lymphocytes to the CNS during EAE induction (40)), suggesting defects in T-lymphocyte migration likely do not fully explain the decreased severity of EAE in HbKO animals. Therefore, in contrast to the *ex vivo* findings, HbKO T-lymphocytes appear to possess many normal functions *in vivo*, at least in the context of EAE.

However, while certain isolated T-lymphocyte functions appeared to be intact, their ability to interact with other cell types essential for EAE development remained unclear. In EAE after CD4^+^ T-lymphocytes are activated and differentiated, T_H_1 and T_H_17 cells can extravasate into the spinal cord and interact with other immune cells, like B-lymphocytes and microglia (41), which are partially responsible for damaging the myelin sheath of the spinal cord and leading to paralysis symptoms. Strikingly, HbKO EAE animals exhibited a significant reduction in plasma IgM and lambda immunoglobulins, two important immunoglobulins in EAE induction (42). Taken together with previous data, this decrease in immunoglobulins suggested an impaired interaction between T-lymphocytes and B-lymphocytes. To preliminarily test this hypothesis, T-lymphocytes were assessed for CD40L and CD27, which are two extracellular markers known to interact with B-lymphocytes (43–47). While no apparent differences were noted in percentage of cells expressing the CD27 receptor in HbKO T-lymphocytes, CD4^+^ HbKO T-lymphocytes did exhibited fewer CD40L^+^ cells after acute activation (measured between its highest expression time point, 4-8 hours after activation (48)). Importantly, the CD40-CD40L interaction has been shown to be critical in EAE development. It has been demonstrated that blocking CD40 throughout the experimental duration significantly blunted EAE symptomology, suggesting the receptor is necessary for demyelination (49). However, additional experiments are required to confirm if the decrease in CD40L expression on HbKO T-lymphocytes is the definitive mechanism underlying the lessened EAE severity.

One potential confounder of the work presented herein is the potential for hemoglobin compensation, particularly of the alpha locus. Due to the importance of Hbα in adult hemoglobin tetramers, many mammalian species (including humans and mice) have evolved two identical copies of the gene: Hbα-a1 and Hbα-a2.

Therefore, since our genetic knock-out model was specific for Hbα-a1, it remains unclear if/how Hbα-a2 plays a role (or compensation) in T-lymphocytes. Anecdotally, we are still able to detect Hbα protein in HbKO T-lymphocytes (data not shown), which suggests Hbα-a2 must be expressed at some level in these cells.

However, the loss of Hbα-a1 (even with potential expression of Hbα-a2) was effective enough to cause a significant phenotype in T-lymphocytes, so compensation from this locus is not fully complete. Additionally, we previously reported detection of beta hemoglobin mRNA in T-lymphocytes as well (8), but found it had very different expression patterns compared to the alpha subtypes. Many hemoglobinopathies are due to mutations or loss of beta hemoglobin loci, but its role in T-lymphocytes remains unknown to date. Last, while we used CD4-cre to induce the recombination of Hbα-a1 in T-lymphocytes, this model initiates knock-out during T-lymphocyte development in this thymus. While we did not observe any baseline changes in total T-lymphocyte numbers or distributions of major subtypes in HbKO animals, it does raise the possibility that the loss of Hbα-a1 during development caused the observed phenotype. Using our Hbα-a1 animals with an inducible T-lymphocyte-driven cre-recombinase in the future will address this potential confounder.

In summary, our data suggest that Hbα is an intricately controlled redox protein temporally expressed in T-lymphocytes subtypes. The loss of Hbα in T-lymphocytes leads to mitochondrial dysfunction, decreased activation potential, and heightened production of inflammatory cytokines, potentially shedding light on the current unknown connection between hemoglobinopathies and autoimmune disorders. Modern molecular tools have allowed us to define the critical nature of Hbα in T-lymphocytes, where it was not even reported to exist, and fully elucidating the regulatory mechanisms and functional role(s) of Hbα within T-lymphocytes will expand our knowledge of T-lymphocyte activation, polarization, and mitochondrial regulation. Additionally, our findings may provide a missing link between autoimmune etiologies in patients with hemoglobinopathies and could pave the way for immune cell-targeted treatments for these patients, rather than solely focusing on erythrocytes.

## Acknowledgements

Funding sources

This work was supported by the National Institutes of Health (NIH) R01HL158521 (AJC) and T32GM135115 (ECR). We thank Dr. Brant Isakson for his contribution of the Hbα-a1 floxed mouse model used in this work. We thank Texas A&M University’s College of Medicine Flow Cytometry and Cell Sorting Facility for the use of their Seahorse Bioanalyzer, as well as Texas A&M University’s Rodent Preclinical Phenotyping Core for the use of the Protein Simple Jess Automated Western Blot and Vetscan HM5 Hematology Analyzer.

## Author contributions

ECR and AJC designed research studies. All authors conducted experiments, acquired data, and/or performed analyses. ECR and AJC wrote the manuscript, while all authors approved the final version of the manuscript. AJC provided funding and experimental oversight.

## Abbreviations

Hbα-a1: hemoglobin alpha-a1 mRNA
Hbα: hemoglobin alpha protein
O_2_: oxygen
NO: nitric oxide
HbKO: T-lymphocyte specific hemoglobin alpha-a1 knock out mice
ROS: reactive oxygen species
EAE: experimental autoimmune encephalomyelitis

---

**Supplementary Figure 1: Generation of T-lymphocyte specific hemoglobin alpha a1 knock-out animals. A**: Schematic of T-lymphocyte specific Hbα-a1 mouse knock out (HbKO) generation. **B**: Results of tail, liver, and isolated pan T-lymphocytes genotyped for Hbα-a1 excision (WT: 1088bp, excised: ∼500bp). **C-F**: Complete blood count results of hemoglobin (**C**), white blood cells (**D**), red blood cells (**E**), and lymphocytes (**F**) from whole blood samples from WT and HbKO animals. **G-H**: Splenic percentages of CD4^+^ (**G**) and CD8^+^ (**H**) T-lymphocytes in WT and HbKO animals. **I-J**: Live cell counts of CD4^+^ (**I**) and CD8^+^ T-lymphocytes (**J**) isolated from whole spleens. **K-L**: Growth curves of activated isolated splenic CD4^+^ (**K**) and CD8^+^ (**L**) T-lymphocytes over 24, 48, 72 hours (N=6). **M-R**: Naïve splenic CD4^+^ and CD8^+^ T-lymphocytes were isolated and measured for MitoSOX (**M-N**), DHE (**O-P**), and DAF-FM (**Q-R**) (mean fluorescence intensity). Statistics (not significant, not shown) were measured using Student’s t-test or two-way ANOVA.

**Supplementary Figure 2: Loss of Hba-a1 in T-lymphocytes does not affect mitochondrial number or mass.** Naïve (A-D) and 72-hour activated (E-H) CD4^+^ and CD8^+^ T-lymphocytes from WT and HbKO animals were assessed for mitochondrial DNA content via qPCR. Mitochondrial mass was assessed in naïve (I) and 72 hour activated (J) T-lymphocytes by determining MFI of MitoTracker Green FM via flow cytometry. Statistics (not significant, not shown) were calculated using two-way ANOVA with Fisher’s LSD multiple comparisons.

**Supplementary Figure 3: Additional cytokines measured in T-lymphocyte time course activation.** Extracellular cytokine protein concentration from activated CD4^+^ (**A-D**) and CD8^+^ T-lymphocytes (**E-H**) from WT and HbKO animals (N=5). Statistics measured using two-way ANOVA.

**Supplementary Figure 4: Fourteen day EAE procedure. A**: Schematic of 14-day EAE experimental design. **B-C**: EAE severity scores (0–5) and mouse weights (g) over 14-day incubation period (WT N=30, HbKO N=25). **D-G**: Protein concentration (pg/mL) of cytokines in plasma. **H-K**: Percentage of CD4^+^ and CD8^+^ present in the spleen and lymph nodes at day 14. **L-O**: Percentage of splenic CD4+ polarized T-lymphocytes. **P-S**: Splenocytes restimulated with 10 μg/mL MOG_35-55_ for 72 hours, then assessed for extracellular cytokine protein concentration (pg/mL per 10^6^ cells). Statistics measured by two-way ANOVA with Šídák’s multiple comparisons test or Student’s t-test where appropriate.

**Supplementary Figure 5: Splenic and spinal cord immune cell percentages do not change between WT and HbKO EAE animals. A-D**: Percentage of inguinal lymph node CD4+ polarized T-lymphocytes. **E-H**: Immune cell populations of B-lymphocytes (**E**), dendritic cells (**F**), granulocytes (**G**) and monocytes (**H**) were assessed by flow cytometry. **I-J**: MitoSOX MFI of CD4^+^ (**I**) and CD8^+^ (**J**) splenic T-lymphocytes. **K**: Percentage of CD45^+^ cells present in the spinal cord. **L**: Live cell count of splenocytes after 72-hour restimulation with 10 µg/mL MOG_35-55_. Statistics (not significant, not shown) were measured using Student’s t-test.

## Notes

### Competing Interest Statement

The authors have declared no competing interest.

### Summary of Updates

1) Revised text regarding hemoglobinopathies throughout document. 2) New supplemental figure

