## Supplementary figures and images for "Hemoglobin alpha regulates T-lymphocyte activation and mitochondrial function"

### Supplemental Figure 1

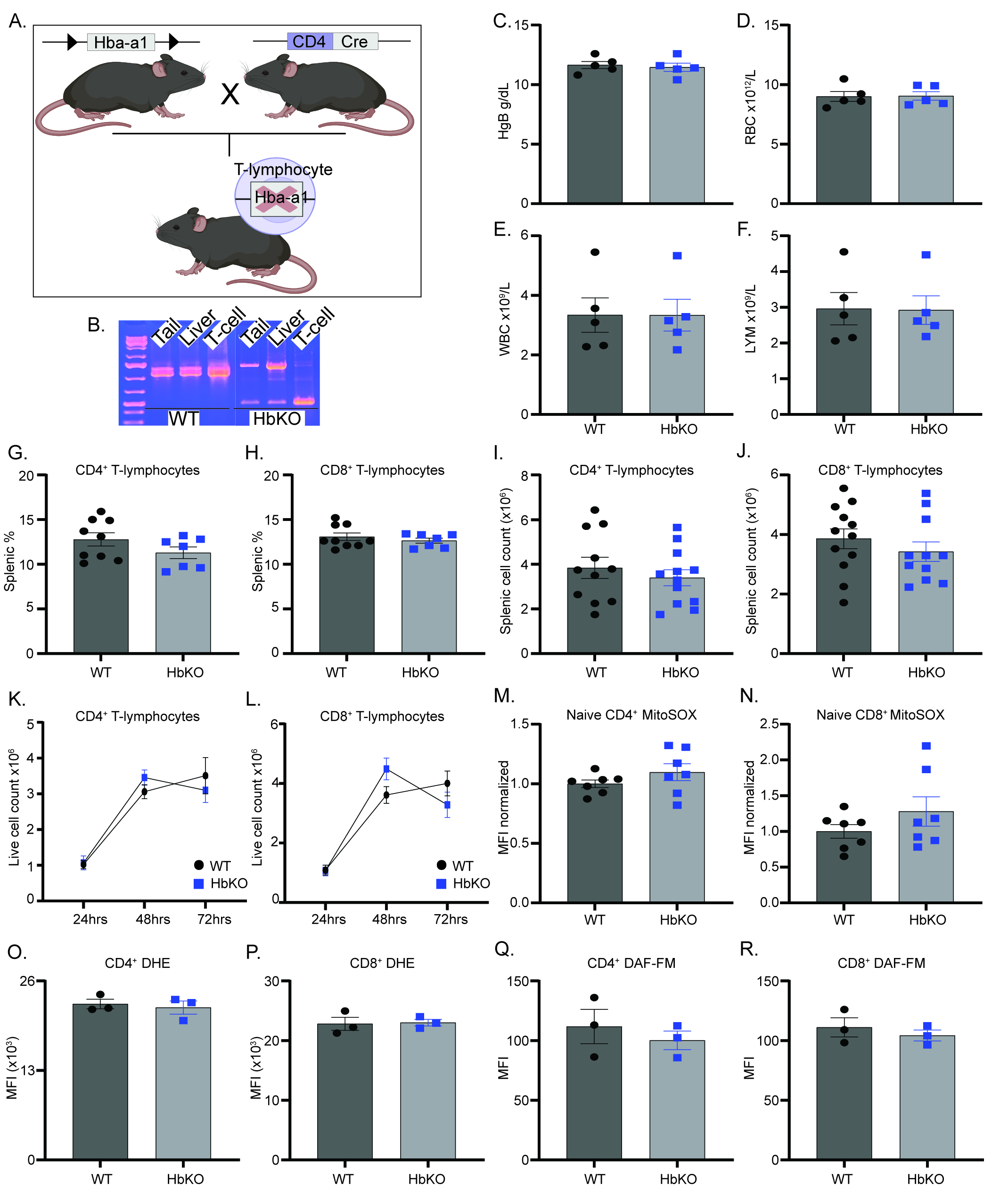

### Supplemental Figure 2

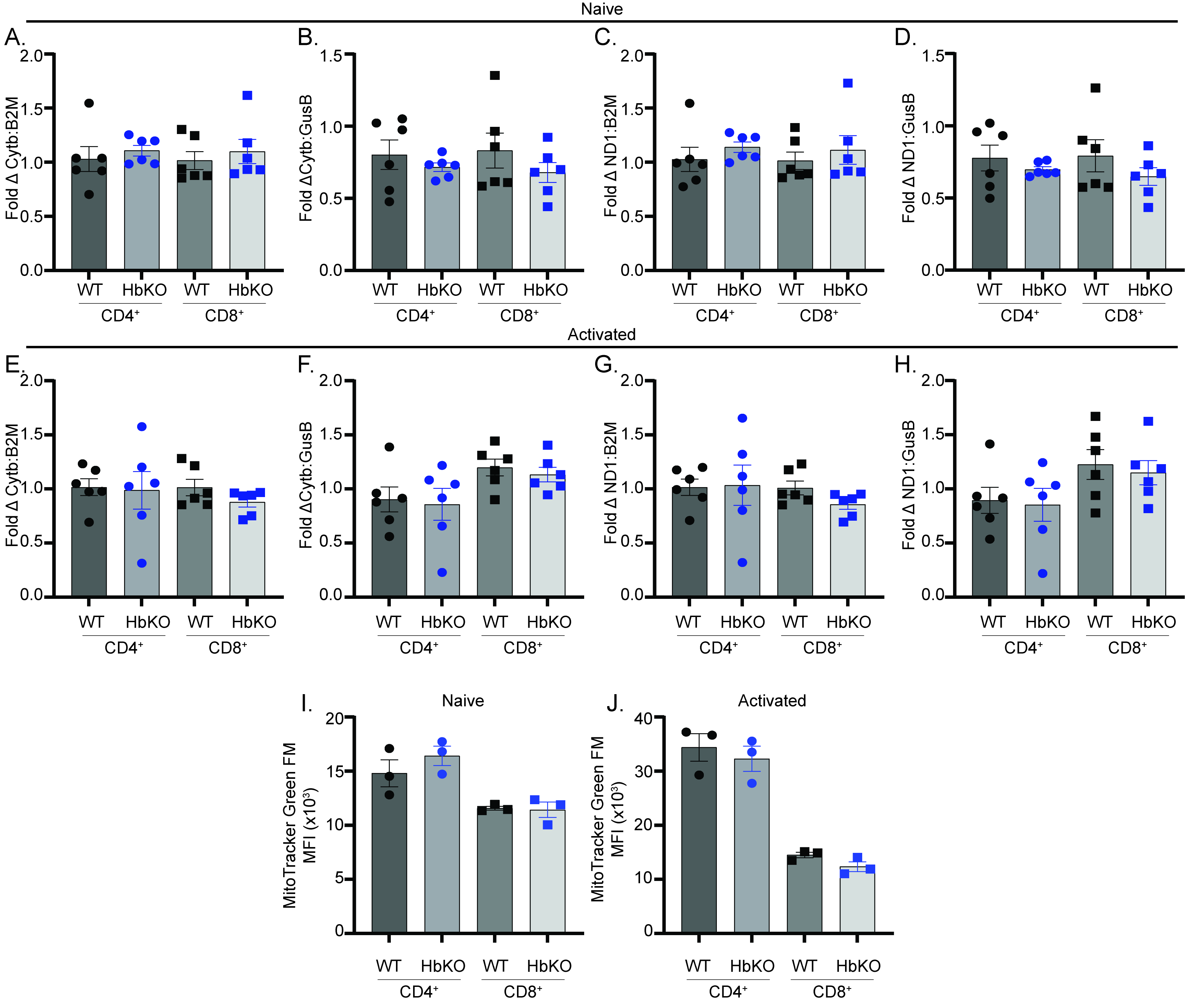

### Supplemental Figure 3

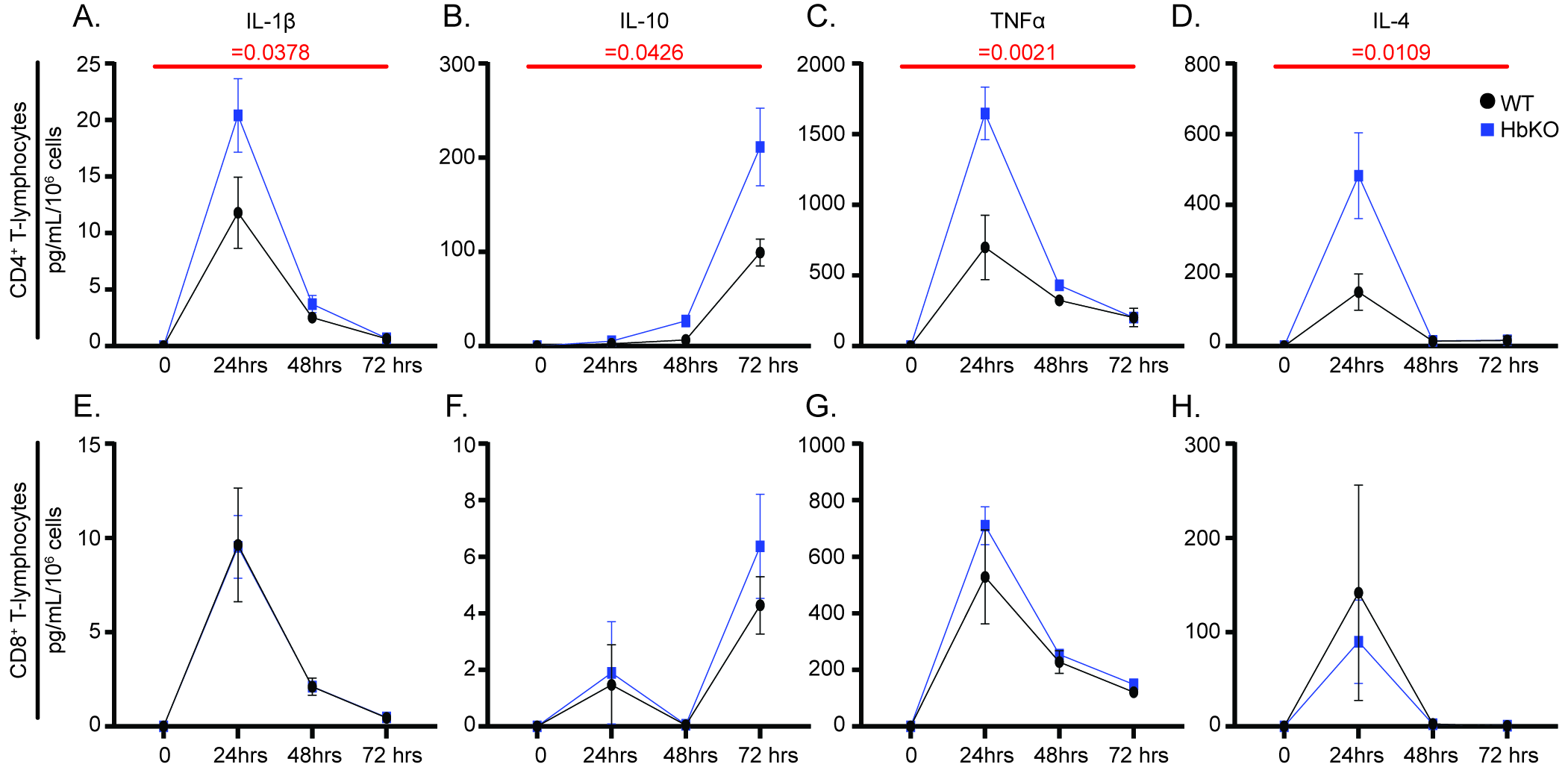

### Supplemental Figure 4

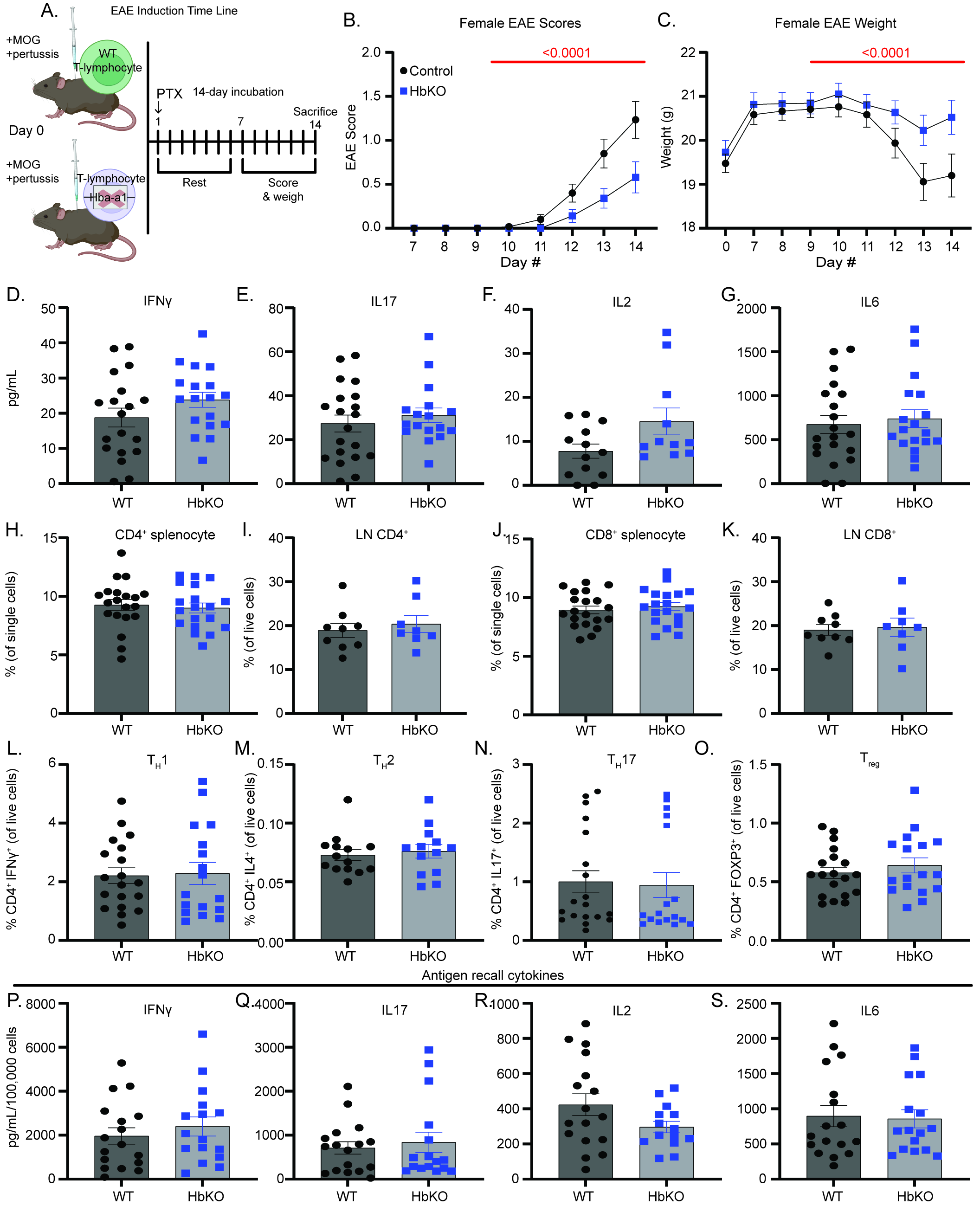

### Supplemental Figure 5

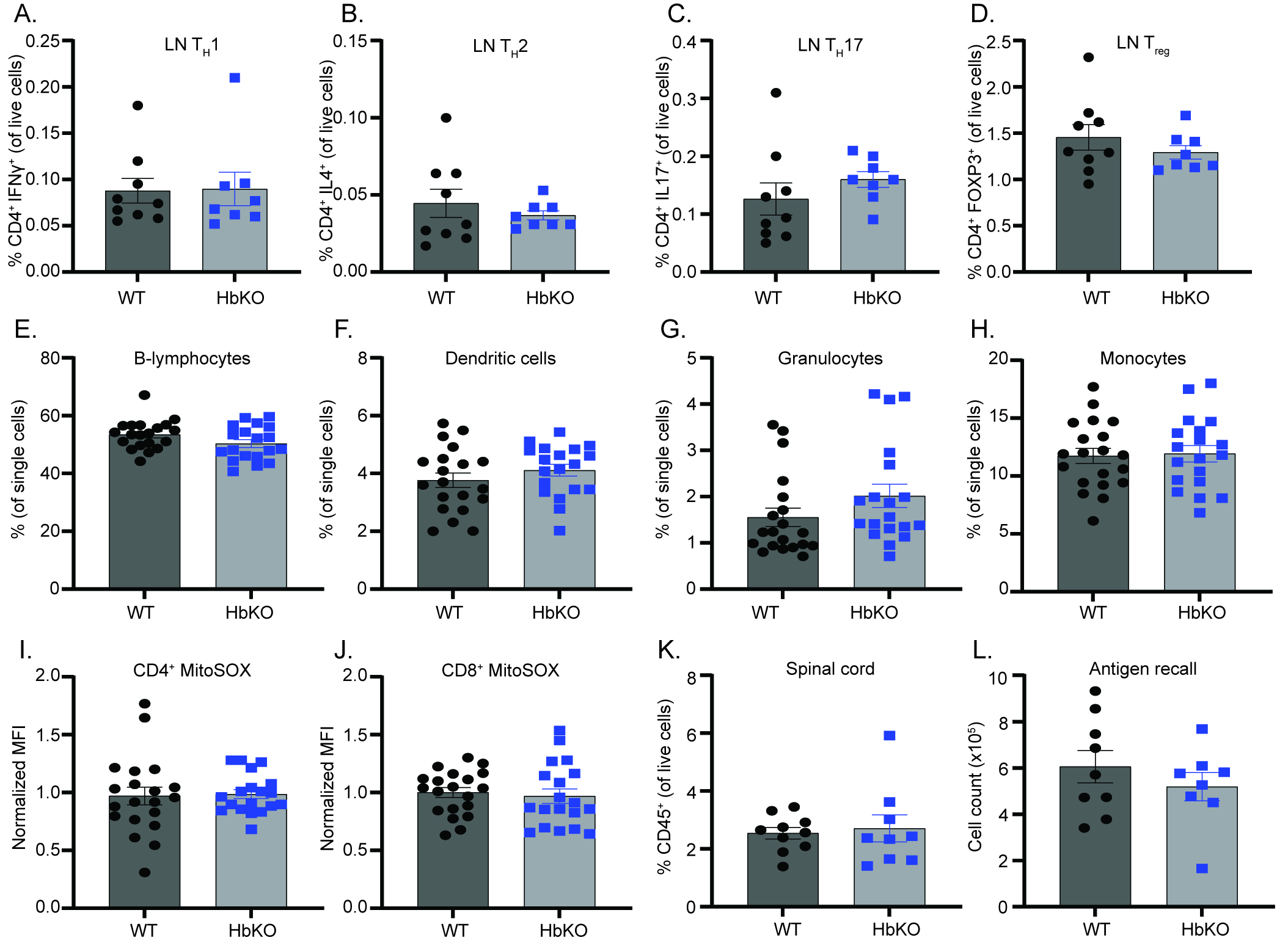
